# Air channels create a directional light signal to regulate hypocotyl phototropism

**DOI:** 10.1101/2023.02.22.529488

**Authors:** Ganesh M. Nawkar, Martina Legris, Anupama Goyal, Emanuel Schmid-Siegert, Jérémy Fleury, Antonio Mucciolo, Damien De Bellis, Andreas Schüler, Christian Fankhauser

## Abstract

In light-limiting conditions, aerial organs of most plants reorient their growth towards the light to improve photosynthesis, through a process known as phototropism^1-3^. The blue light receptors phototropin control phototropic responses through light-induced protein kinase activity^4^. Current models posit that asymmetric activation of these sensory receptors across a unilaterally illuminated organ leads to asymmetric distribution of the growth hormone auxin ultimately leading to growth re-orientation^4,5^. However, the tissue properties required to generate a light gradient across the stem triggering phototropism remain unclear^1^. Here we show that inter-cellular air channels^6,7^ are required for an efficient phototropic response. These channels enhance light scattering (refraction and reflection) in Arabidopsis hypocotyls thereby enhancing the light gradient across the photo-stimulated organ. We identify an embryonically expressed ABC transporter that is required to keep air in inter-cellular spaces in seedlings and for efficient phototropism. Our work suggests that this transporter shapes cell wall properties to maintain air between cells. Moreover, we establish the functional importance of inter-cellular air channels in the hypocotyl for phototropism.

## Main text

In flowering plants light direction is sensed by the blue-light (BL) absorbing phototropin photoreceptors (phot1 and phot2 in Arabidopsis)^5^. This typically leads to growth towards the light or positive phototropism in aerial organs such as hypocotyls and stems^4^. This response is believed to contribute to the optimization of photosynthesis particularly in limiting light conditions^2,3,8^. In dicotyledons like *Arabidopsis thaliana*, the upper hypocotyl is both the site of light perception and the site of differential growth ultimately leading to organ repositioning^9,10^. Upon unilateral BL irradiation differential phot activation between the lit and the shaded side of the seedling is considered as the first step triggering phototropism^5,11^. Substantial progress was made in elucidating the downstream steps linking phot activation and the differential growth response^5^. However, the optical features of light-sensing tissues enabling the formation of a light gradient that are required for a phototropic response remain poorly characterized^1^.

### Transparency of hypocotyls causes phototropic defects

In a screen for Arabidopsis seedlings with reduced phototropism we identified a mutant with transparent hypocotyls (Fig. 1). The causal gene was mapped to *ATP-BINDING CASETTE G5* (*ABCG5*), which was confirmed by comparing the phenotype of multiple alleles and by complementation (Extended data Fig. 1). We hypothesized that the phototropic defect in the mutant was caused by enhanced light transmission and a shallower light gradient in the upper part of the hypocotyl. To test this hypothesis, we compared *abcg5-5* (from now on *abcg5*) and two previously identified *cristal* mutants (*cri7* and *cri8*), which also have transparent hypocotyls^12^. The defective gene in these mutants is not known^12^, but they are not allelic^12^ and we found that the *ABCG5* gene was unaltered (see Supplementary Methods). The three mutants had similarly enhanced light transmission in the hypocotyl (Fig. 1a). In response to low unilateral BL *abcg5, cri7* and *cri8* growth orientation was random, while the wild type (WT) aligned with the light source and *phot1* was unresponsive (Fig. 1b). To determine whether these mutants have a specific hypocotyl tropism defect, we analyzed their gravitropic response, which also relies on asymmetric growth caused by redistribution of the growth hormone auxin ^13^. An auxin transporter mutant deficient in three *PIN-FORMED* genes (*pin3pin4pin7*) showed the expected inability to re-orient its hypocotyl upon gravistimulation (Fig. 1c). In contrast, the response of *abcg5* and *cri7* was similar to the WT, while *cri8* showed a reduced response to gravity (Fig. 1c). Both phototropism and gravitropism depend on growth; we therefore measured hypocotyl elongation during the phototropic experiment and found that while *abcg5* hypocotyls grew like the WT, *cristal* mutants showed a growth defect (Extended data Fig. 2). We conclude that having a transparent hypocotyl correlates with an altered ability to respond to light direction and pursued our study with the *abcg5* mutant due to the pleiotropic nature of *cristal* mutants^12^.

**Figure 1.**
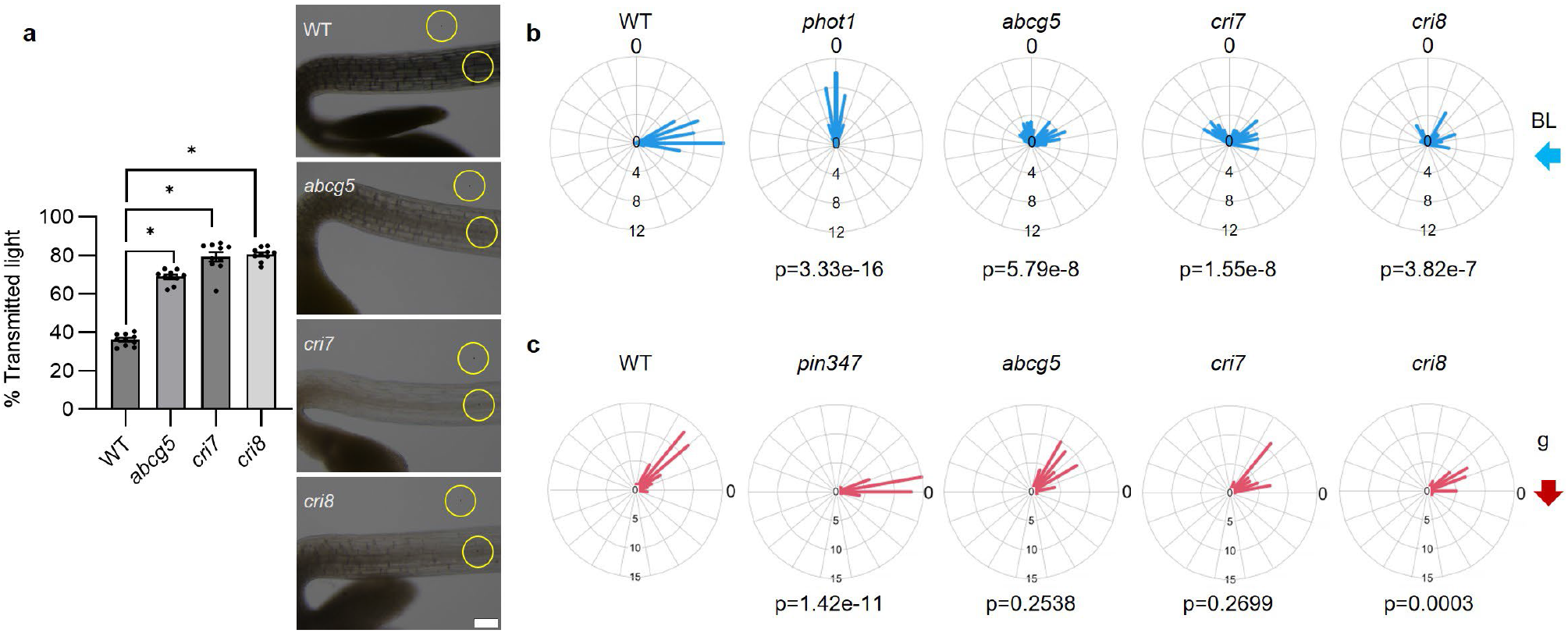
Transparent hypocotyl mutants are specifically impaired in the perception of light direction. (a) Left: Hypocotyl transmittance quantified from bright field microscopy images of 3-day-old etiolated seedlings. Right: representative images used for quantification. Scale bar 100 μm. Transmittance was quantified as the ratio between the mean gray value of a circular region of interest (ROI) on the hypocotyl and a ROI in an empty region of the image (yellow circles). Data are mean ± SEM of n=8-9 seedlings. * P<0.0001 in ANOVA followed by Dunnett’s multiple comparisons test. (b) Radar plots showing the results of phototropic assays. 3-day-old etiolated seedlings growing on vertical plates were treated with unilateral blue light (0.025 μmol m^-2^ s^-1^) for 24 h and final growth angle relative to vertical was measured. Blue bars’ length represent the relative frequency of bending angles in 10° intervals. Data distribution for each genotype was compared to the WT using a Kolmogorov-Smirnov test (P-values below each graph). (c) Radar plots showing the results of gravitropic assays. 3-day-old etiolated seedlings growing on vertical plates were rotated by 90° and growth angle relative to horizontal was measured 24h later. Red bars’ length represent the relative frequency of bending angles in 10° intervals. Data distribution for each genotype was compared to the WT using a Kolmogorov-Smirnov test (P-values below each graph).

If the *abcg5* phototropic defect were due to impaired light direction sensing, we would expect that it requires active phototropins. Consistent with this hypothesis, the *phot1abcg5* double mutant behaved like a *phot1* mutant in response to low BL (Extended data Fig. 3a), while the *abcg5* mutants showed many seedlings growing in the opposite direction (Fig. 1, Extended data Fig. 3a). These experiments were performed with seedlings growing on vertical plates in contact with the media. Given that agar scatters light and that plants were growing at the media-air interface, the light environment in this experimental setup was probably relatively complex. Thus, the following experiments were performed in free standing etiolated seedlings to obtain a simpler directional light cue (see Supplementary Methods). To better characterize the phototropic phenotype of the *abcg5* mutant, we analyzed pulse-induced first positive phototropism, which corresponds to the conditions where light fluence (μmol m^-2^) is proportional to the phototropic response^14^. We irradiated etiolated seedlings with a 1-minute BL pulse of various intensities (1.7, 0.17, 0.017, and 0.0017 μmol m^-2^ s^-1^). In agreement with previous reports^15,16^, WT plants showed a bell-shaped fluence-response curve, whereas *abcg5* mutants showed a severe phototropic impairment (Extended data Fig. 3b). Next, we examined time-dependent phototropism^14^, by irradiating etiolated seedlings continuously with different BL fluences (0.025, 0.125, and 2.5 μmol m^-2^ s^-1^) and recorded growth re-orientation over 6 hours. We found that WT plants took longer to reach maximum curvature with increasing light intensity, confirming earlier studies^16^. Strikingly, the *abcg5* mutant took much longer than the WT to reach maximum curvature at all tested light intensities (Extended data Fig. 3c). We used a bi-directional light treatment to further test the ability of seedlings to respond to complex light environments. In such conditions WT plants grow towards the stronger light source^1^. In our test area seedlings received a similar light intensity but different light gradients depending on their position (P) (steep (P1, P6), medium (P2, P5) and shallow (P3, P4)) (Extended data Fig. 3d). The ability of *abcg5* mutants to grow towards stronger light was reduced particularly when the gradient was shallow (Extended data Fig. 3e). Moreover, their response was slower in this situation (Extended data Fig. 3f). Collectively our phototropism experiments indicated that *abcg5* mutants showed a phototropic defect in all tested conditions. The phenotype was particularly pronounced in response to very low light fluences, high fluence rates, and in complex light environments as on the surface of plates or with bi-directional irradiation.

Despite these obvious phototropic defects, we found that phot1-mediated phosphorylation events occurring within minutes of light perception were not altered in the *abcg5* mutant (Extended data Fig. 4). Indeed, the BL-induced mobility shifts of phot1 and its targets NPH3 and PKS4 observed on SDS-PAGE gels were normal in *abcg5*. This is consistent with the *abcg5* phototropic defect being due to a reduced ability to establish a light gradient across the hypocotyl rather than in downstream phot signaling (Extended data Fig. 4)^17-19^. Moreover, the defective BL response of *abcg5* was specific to phototropism as BL-induced inhibition of hypocotyl elongation was unaltered in *abcg5* (Extended data Fig. 5a). Transparency of the *abcg5* mutant was restricted to the embryonic phase, while true leaves and other plant organs developed similarly to the WT (Extended data Fig. 5b). This allowed us to test the phototropic response in petioles. Interestingly, *abcg5* mutant petioles showed a WT response (Extended data Fig. 5c), hence we conclude that the phototropic defect of the mutant is restricted to transparent organs. Analysis of *ABCG5* gene expression showed that the gene was particularly strongly expressed in developing embryos, with strongest expression in the cortex (Extended data Fig. 5d), while we did not observe expression with a reporter line in seedlings. Of note, while we could complement the mutant by expressing the *GFP-ABCG5* transgene from the *ABCG5* promoter, complementation was unsuccessful when the construct was driven by the viral *35S* promoter (Extended data Fig. 1c). Given that the *35S* promoter does not drive gene expression during the early stages of embryogenesis in other species^20,21^, our complementation assays suggest that embryonic expression of *ABCG5* is functionally important. This may also explain the seedling-specific phenotype of *abcg5* mutants, while the mutant showed no obvious phenotype later in development (Extended data Fig. 5). We conclude that *abcg5* has a specific phototropic defect in seedlings that is most likely due to hypocotyl transparency.

### ABCG5 is required for inter-cellular air channels formation

Light absorbing pigments were proposed to contribute to light gradient formation across photo-stimulated plant tissues thereby enabling phototropism ^22,23^. We therefore analyzed the absorption spectrum of soluble crude extracts from etiolated seedlings and found that *abcg5* extracts showed an absorption spectrum comparable to the WT (Extended data Fig. 6). A recent study showed that ABCG5 is required for cuticle development in cotyledons with the mutant showing higher permeability of cotyledons in light-grown seedlings^24^. Thus, we evaluated the hypothesis that a defect in cuticle development in the hypocotyl explains the phototropic defect and hypocotyl transparency in etiolated seedlings. First, we assessed the cuticle structure by transmission electron microscopy (TEM). We did not find large differences between WT and *abcg5*, although as reported for the cotyledons, the cuticle in *abcg5* was slightly less compact than in the WT (Extended data Fig. 7a). However it was still functional, since the toluidine blue cuticle permeability test showed that *abcg5* and WT had a similar permeability, which contrasted with a severe cuticular deficient mutant (long-chain acyl-CoA synthetase2, *lacs2*) ^25^, (Extended data Fig. 7b). Moreover, despite cuticular defects, hypocotyls of *lacs2* mutants were not transparent and had a robust phototropic response indicating that hypocotyl transparency and the phototropic defect in *abcg5* are presumably unrelated to the cuticle (Extended data Fig. 7c). Light-grown *abcg5* seedlings sink in water suggesting that they contain less air than the WT^24^. We performed a floating assay with dark-grown seedlings and found that dissected roots and hypocotyls as well as whole seedlings of the *abcg5* mutant sank more frequently than the WT, indicative of reduced air content in the *abcg5* mutant (Fig. 2a). Air channels were previously observed in embryos and hypocotyls of several species^6,7^. They are present at the tricellular junctions between cortex cells or between cortex and epidermal cells^6^. Thus, we analyzed transverse cuts of etiolated hypocotyls using cryo-scanning electron microscopy (cryo-SEM) and TEM. Both in WT and *abcg5* hypocotyls we observed intercellular spaces at the tricellular junctions formed by epidermal and cortex cells. However, those spaces looked empty in the WT, while they were filled in the *abcg5* mutant (Fig. 2b, Fig. 2c, Extended data Fig. 8). Moreover, in TEM images the WT showed a well-defined electron dense layer in the outer side of the cell wall surrounding the intercellular spaces. Interestingly, in *abcg5* mutants this layer was diffuse, heterogeneous, and sometimes absent (Fig. 2b, Extended data Fig. 8). This structure may correspond to the “splitting layer”^26^, that was hypothesized to contain hydrophobic material sealing the intercellular space and allowing the formation of air channels. To further investigate whether the difference between WT and *abcg5* was the presence of air in intercellular spaces, we used 3D-non-destructive X-ray microtomography^7,27^. We detected air channels in the longitudinal direction in the WT but not in *abcg5* mutants (Fig. 2d). Collectively, our data suggest that the difference in light transmission between the WT and *abcg5* may be explained by the presence of air in the intercellular spaces of the WT but not in the mutant.

**Figure 2.**
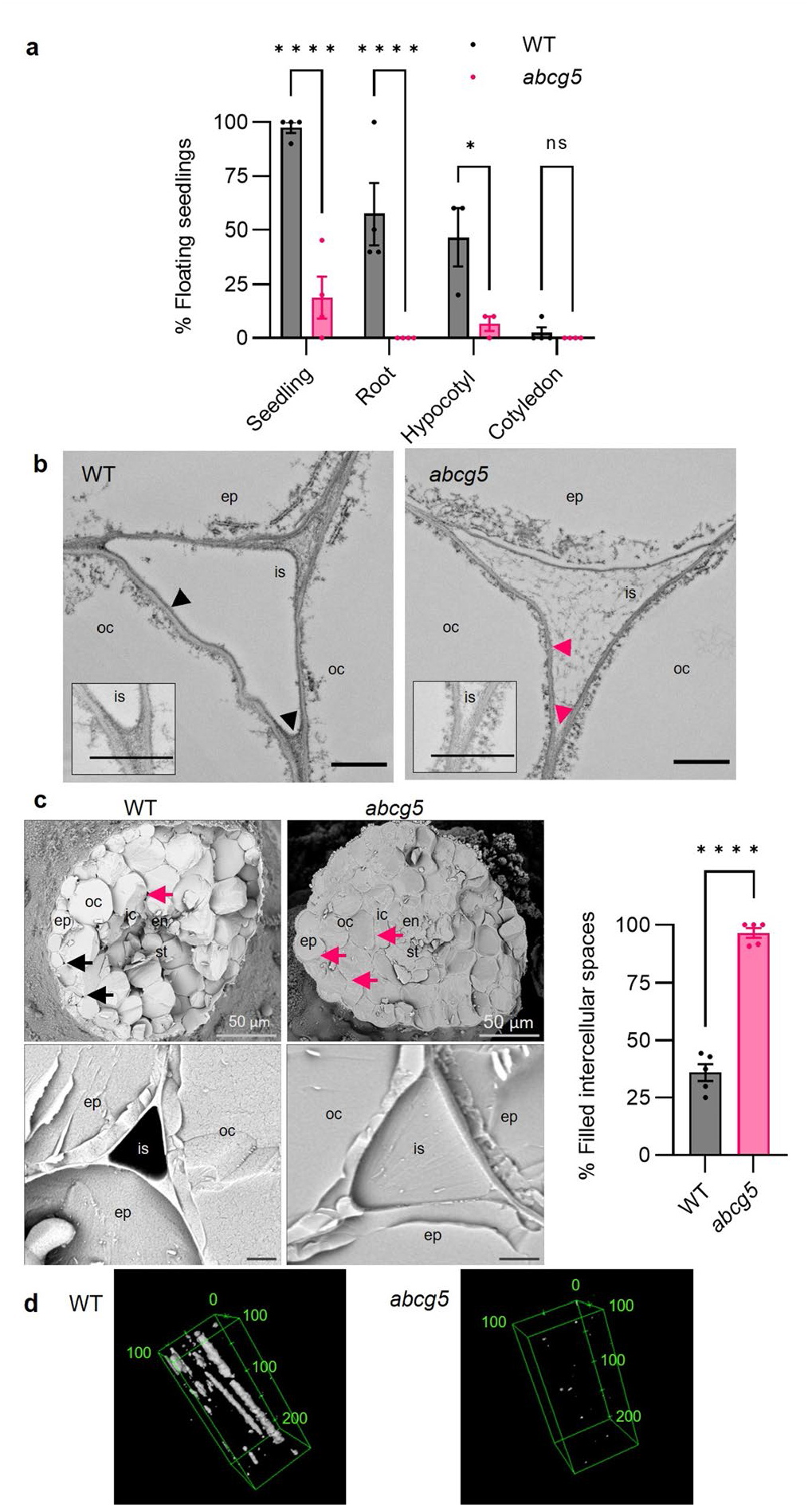
The difference between WT and mutant is the presence of air bubbles. (a) Floating assay of 3-day-old etiolated seedlings, or dissected root, hypocotyl, and cotyledons of WT and *abcg5*. Data are mean ± SEM, n=4. *P<0.05, ****P<0.0001, and NS, no significant in an ANOVA followed by Šídák’s multiple comparisons test. (b) Transmission Electron Microscopy images of intercellular spaces (is) in WT and *abcg5*. Inset: close-up look of a corner. Scale bar 1 μm. Black arrowheads mark the electron-dense layer in the WT and pink arrowheads mark similar positions in *abcg5*. (c) Left: Cryo-scanning electron microscopy images of freeze fractured transverse cuts of 3-day-old etiolated hypocotyls (top) or close-up looks of intercellular spaces (bottom, Scale bar 1 μm). Black arrows mark empty intercellular spaces. Pink arrows mark filled intercellular spaces. ep, epidermis; oc, outer cortex; ic, inner cortex; en, endodermis; st, stele; is, intercellular space. Right: Quantification of images. Data are mean ± SEM. n=5 hypocotyls. ****P<0.0001 in an unpaired t-test. (D) 3D representations of X-ray microtomography images of WT and *abcg5* hypocotyls. Black: background. White: air. Green lines show the analyzed volume. Numbers in green are dimensions in micrometers.

### Air channels contribute to the formation of directional light cues

To better characterize the optical properties of etiolated seedlings we used an integrating sphere allowing measurements of total transmittance and reflectance along with light scattering, i.e., diffused transmittance and reflectance. As observed with our light microscopy measurements (Fig. 1), *abcg5* seedlings transmitted more light than the WT (Fig. 3a). Moreover, we filled the air spaces in WT seedlings by water infiltration and found that infiltrated WT samples showed optical properties similar to the *abcg5* mutant. In contrast, infiltration of *abcg5* samples did not lead to any significant changes in optical properties (Fig. 3a). Our data showed that *abcg5* mutants and infiltrated seedlings showed reduced diffused transmitted light, reflected light and diffused reflected light (Fig. 3a). This data is consistent with air channels enhancing light scattering in plant tissues due to the strong difference in the refractive index of air compared to water, cellular fluid and cell walls^28^. To understand how changes in optical properties affect the light microenvironment within the hypocotyl, we used confocal microscopy. We reconstructed hypocotyl transverse cuts using Z-stacks of transmitted light images. These images showed a similar pattern in all samples, but the contrast was higher in the WT compared to the *abcg5, cristal* mutants and infiltrated samples indicating stronger light scattering in the WT (Fig. 3b, Extended data Fig. 9a). Moreover, combining these images with images of membrane-associated fluorescent proteins we could determine that the areas of strong scattering coincided with the intercellular spaces at tricellular junctions (Fig. 3c, Extended data Fig. 9b,c). Collectively, our optical characterization of etiolated hypocotyls showed that intercellular air channels contribute to light scattering thereby limiting light transmittance across the hypocotyl.

**Figure 3.**
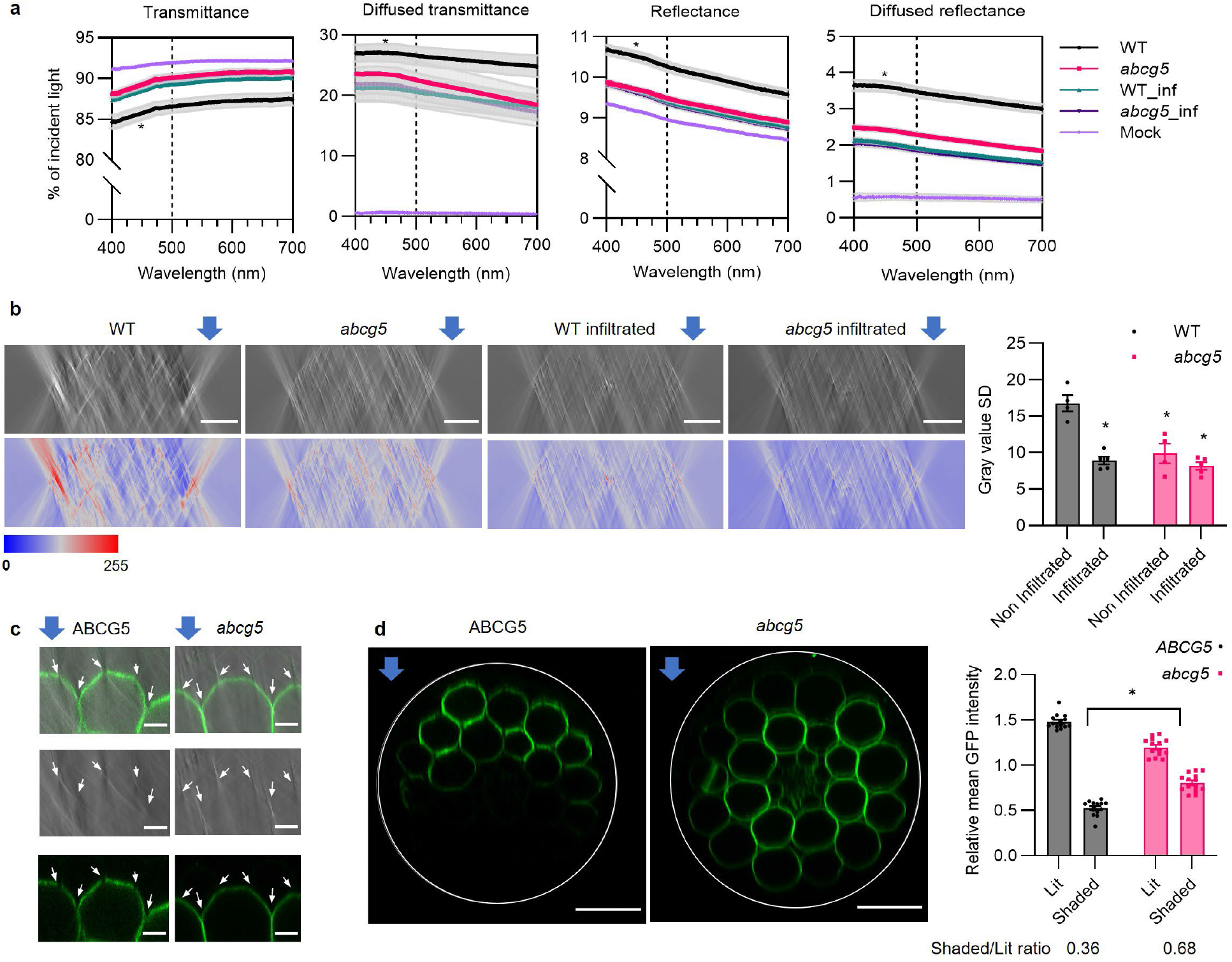
Air channels create a directional light signal. (a) Optical properties of seedlings measured with the integrating sphere. Data are mean ± SEM of 5-8 samples from two experiments. Treatments were compared calculating the area under the curve in the blue range of the spectrum (400-500nm). ANOVA followed by Tukey’s multiple comparisons test showed significant differences between WT and all other treatments (*P<0.001). (b, c, d) xz orthogonal views of bright field and/or fluorescence confocal images of etiolated hypocotyls. Blue arrows indicate the direction of the incident light. (b) Left: Representative images in greyscale (upper row) or using an artificial LUT (lower row). Scale bar: 50 μm. Right: Quantification of images. Data are mean ± SEM of five (non-infiltrated) or six (infiltrated) samples. * P<0.001 in an ANOVA followed by Sidak’s multiple comparisons test vs. WT non-infiltrated. (c) Etiolated hypocotyls expressing *pPHOT1:PHOT1-GFP* (green) in an ABCG5 wild type (*ABCG5*) or mutant (*abcg5*) backgrounds. Top: Merge, Middle: Bright field, Bottom: Fluorescence. White arrows show the intercellular spaces between epidermal and cortex cells. Scale bar: 10 μm. (d) Left: Fluorescence images of etiolated hypocotyls expressing *pPHOT1:PHOT1-GFP* (green) in an ABCG5 wild type (*ABCG5*) or mutant (*abcg5*) backgrounds. The white circle shows the limits of the hypocotyl epidermis. Scale bar: 50 μm. Right: Quantification of images. For each image, the mean grey value of the whole transverse cut, as well as of the half closer to the light source (lit) and farther from it (shaded) were quantified. Plotted values are mean grey values of each side relative to the mean grey value of the whole image. The shaded/lit ratio is calculated from these relative values. For each genotype data was fitted to a linear regression and the slopes were compared using a t-test showing significant differences between WT and *abcg5* (* P<0.05). n = 14

To visualize the light gradient across an etiolated hypocotyl we used a *pPHOT1:PHOT1-GFP* line either in *phot1phot2* ^29^ (*ABCG5*) or in a *phot1phot2abcg5* (*abcg5*) background. We made Z stacks across the entire width of the hypocotyl using a confocal microscope equipped with a BL laser. As observed previously^9,29^, the GFP signal was strongest in cortex cells due to the *pPHOT1* expression pattern (Fig. 3c,d). In the *ABCG5* background we noticed GFP signal gaps along the plasma-membrane, which correspond to the position of intercellular air spaces formed between the epidermis and cortex or cortex-cortex junctions (Fig. 3c, Extended data Fig. 9b, c). These fluorescence gaps were not observed in the *abcg5* background (Fig. 3c, d, Extended data Fig. 9b). Similar gaps were observed when using a line expressing 35S promoter-driven plasma-membrane associated GFP (35S:myri-GFP) in the WT, indicating that the effect of air channels on fluorescence visualization is not specific to PHOT1 (Extended data Fig. 9c). To determine the effect of light scattering on the light gradient across the hypocotyl, we compared the PHOT1-GFP fluorescent signal on the lit versus shaded sides of the hypocotyl (Fig. 3d). Our quantifications showed that the light gradient was significantly steeper in the WT than in the *abcg5* mutant or infiltrated seedlings (Fig. 3d, Extended data Fig. 9d). Taken together our results show that air channels present in the WT enhance light scattering thereby leading to a stronger light gradient across a unilaterally illuminated hypocotyl. The characterization of the *abcg5* mutant, which lacks these air channels, shows that this is functionally relevant for the detection of directional light cues.

## Conclusion

The photoproduct-gradient model of phototropism states that the difference in the levels of photoproduct between the lit and shaded side regulates phototropic bending^1^. Some experimental evidence supports the importance of light absorbing pigments in the establishment of a light gradient^22,30^. However, it was also noted that light scattering (in the sense of light diffusion by refraction and reflection) is likely to be important given that the light path inside plant tissues traverses media with different refractive indices (RI, air RI= 1, cell wall RI= 1.42, and cellular fluid RI=1.33)^28^. Our data using the *abcg5* mutant and water-infiltrated WT hypocotyls is consistent with earlier work in other species in showing that air channels in hypocotyls contribute to such a light gradient (Fig. 3)^28^. The shallower gradient in *abcg5* presumably explains why the mutant has more difficulty growing towards the light maximum in complex light environments and shows a delay in responding to unilateral light (Fig. 1, Extended data Fig. 3). The residual phototropic response in the mutant can be explained by the presence of a shallow gradient (Fig. 3). This gradient is presumably due to the difference in refractive indices between cellular fluids and the cell wall, as it could be largely eliminated by infiltrating sunflower hypocotyls with cedarwood oil, which created a medium with a homogeneous RI^28^. Our work shows that ABCG5 is important to seal air channels which were reported in the hypocotyl of several species^6,7,26,27^. These channels form following the separation of a poorly characterized cell wall layer called the “splitting layer”^26^. How this process occurs is poorly understood but interestingly it was proposed that this layer comprises a lipophilic film analogous to suberin or cutin^26^. ABCG5 is an ABC transporter related to transporters implicated in delivering precursors of the cuticular layer to the extracellular space ^31^. Strong expression of *ABCG5* in embryonic cortex cells (Extended data Fig. 5d) and its localization at the plasma-membrane^24^ suggest that it may be implicated in the deposition of lipidic precursors required to seal air channels in embryonic tissues. Consistent with this idea, we described an electron-dense layer visible by TEM surrounding the intercellular space in the WT, which is reduced or absent in *abcg5* mutants (Fig. 2b, Extended data Fig. 8). Finally, our work shows that these air channels which play a role in gas exchange ^6,7,27^ also shape the optical properties of translucent tissues enabling them to provide directional light information to plants.

## Acknowledgements

We thank Prof. Sara Simonini and Dr. Célia Baroux for their help imaging embryos and for fruitful discussions. The X-ray microtomography experiments were done at the EPFL Platform for X-Ray Radioscopy and Tomography (EPFL PIXE) with the help of Albert Taureg. Confocal microscopy was done at the Cellular Imaging Facility (CIF) at the University of Lausanne. The identification of ABCG5 as the causal mutation of the transparent hypocotyl was done at the Genomic Technologies Facility (GTF) at the University of Lausanne. Dr. Johanna Krahmer developed the script to plot phototropism and gravitropism data. This work was supported by the University of Lausanne, the Swiss National Science Foundation (grants no. 310030B_179558 and 310030_200318 to C.F.), Human Frontiers Science Program (LT000829/2018-L to M.L.), European Commission Marie Curie fellowship (grant no. H2020–MSCA–IF–2017–796443 to M.L. and grant no. H2020-MSCA-IF-2018-843247 to G.M.N).

## Author contributions

G.M.N, M.L., A.G. and C.F. conceived the original research plans. G.M.N, M.L. and A.G. performed the experiments and analyzed the data. E.S. collaborated with data analysis for the identification of the causal mutation in *abcg5-5*. J.F and A.S. collaborated with the integrating sphere measurements and with discussions about optics. A.M. and D.D.B. performed the Cryo-SEM and TEM experiments, respectively. G.M.N., M.L. and C.F. wrote the article with contributions of all the authors.

## Competing interests

The authors declare no competing interests.

---

**Extended data Figure 1.**
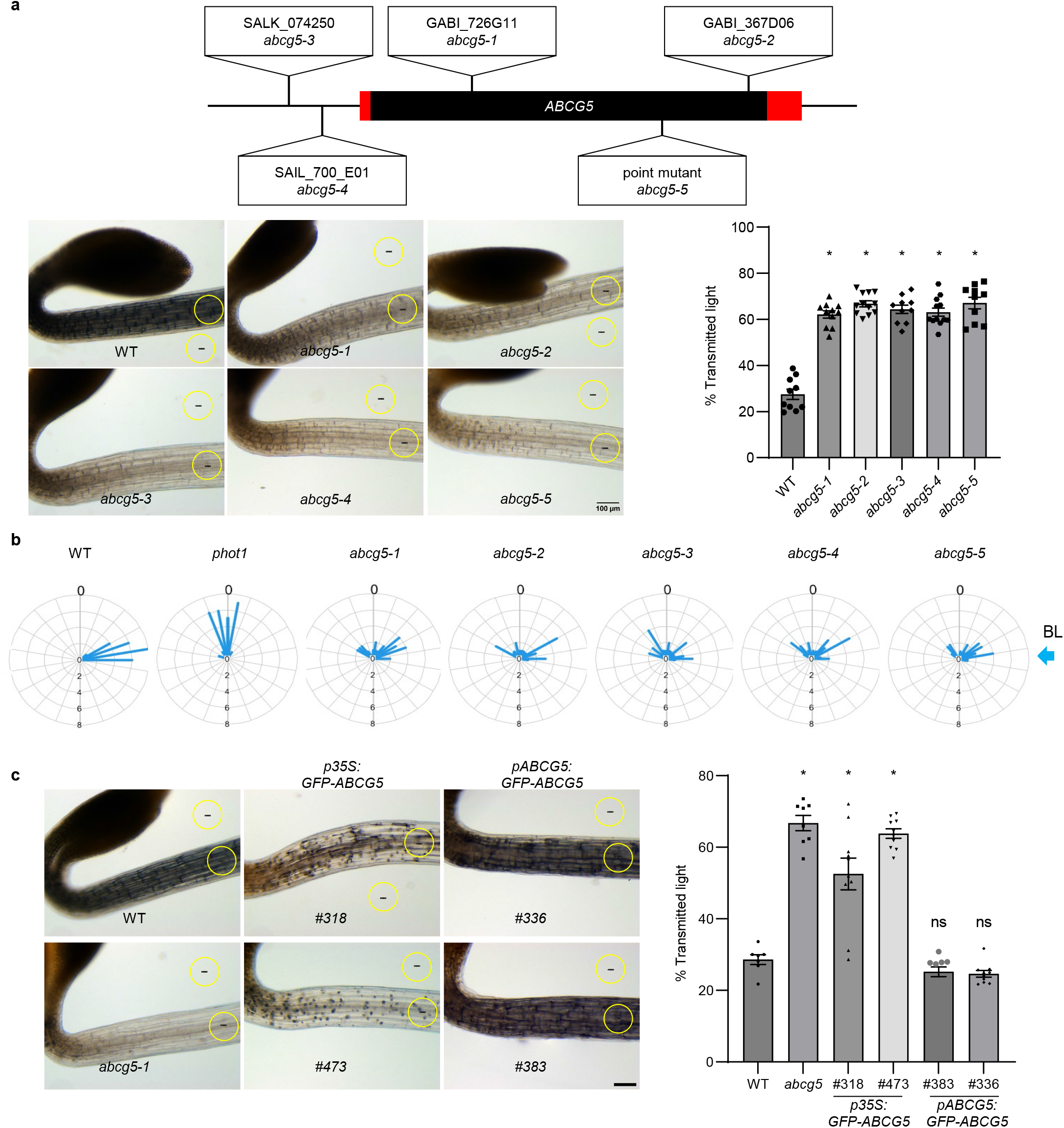
ABCG5 is required for normal hypocotyl development and phototropism. (a) Top: mic structure of ABCG5. The exon and UTRs are shown in black and red color, respectively. The T-DNA insertion in the exon and promoter region are present in lines *abcg5-1, abcg5-2, abcg5-3*, and *abcg5-4*, respectively. The point mutation at G538R obtained from the EMS screen is the *abcg5-5* allele. Bottom left: Representative transmission images of 3-day-old etiolated seedlings. Bottom right: transmitted light calculated measuring a ratio of the gray values in the yellow circular region of interest (ROI) on the hypocotyl and empty ns. (mean ± SEM; n=8-10 seedlings, *P<0.0001, ns not significant in an ANOVA followed by Dunnett’s multiple comparisons test. (b) Radar plots showing the random bending of *abcg5* alleles. Three-day-old etiolated ings grown on vertical plates were treated with unilateral blue light (0.025 μmol m^-2^ s^-1^) for 24 h and measured final growth angle relative to vertical. The bar length in the radar plot represents the relative frequencies of hypocotyl angles in 10 ° intervals. n=24-28. (c) Left: Representative transmission images of 3-day-old etiolated *abcg5* complementation lines. Stable T3 transgenic lines in the *abcg5* background were generated by expressing GFP-ABCG5 under *35S* (#318, #418) and *ABCG5* (#336, #383) promoters. Right: quantification as in (a).

**Extended data Figure 2.**
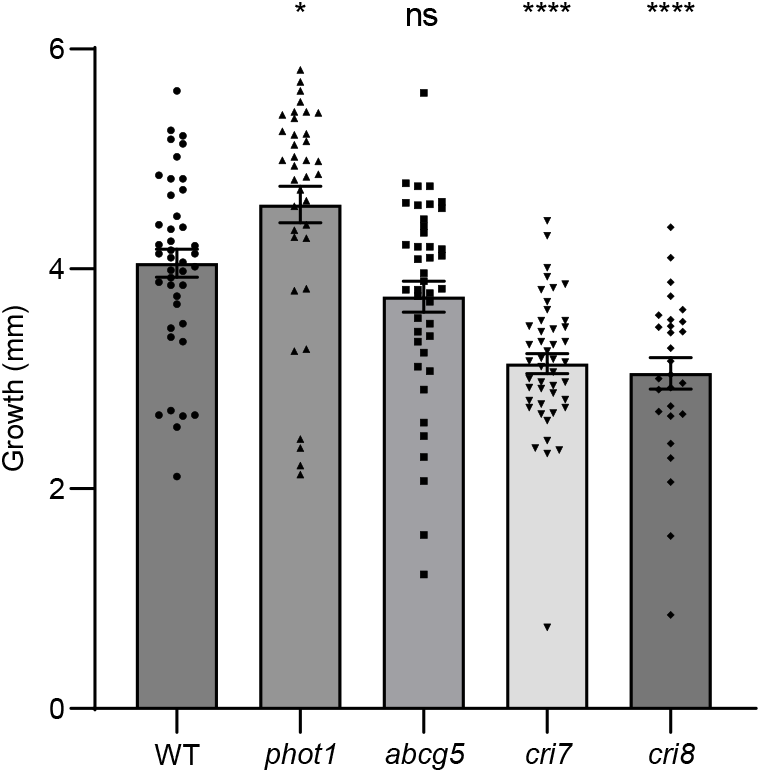
Hypocotyl growth is impaired in *cri* mutants. Hypocotyl elongation during 24h. Data are mean ± SEM of n=27-47 seedlings. *P<0.05, ****P<0.0001, and NS, no significant in an ANOVA followed by Dunnett’s multiple comparisons test.

**Extended data Figure 3.**
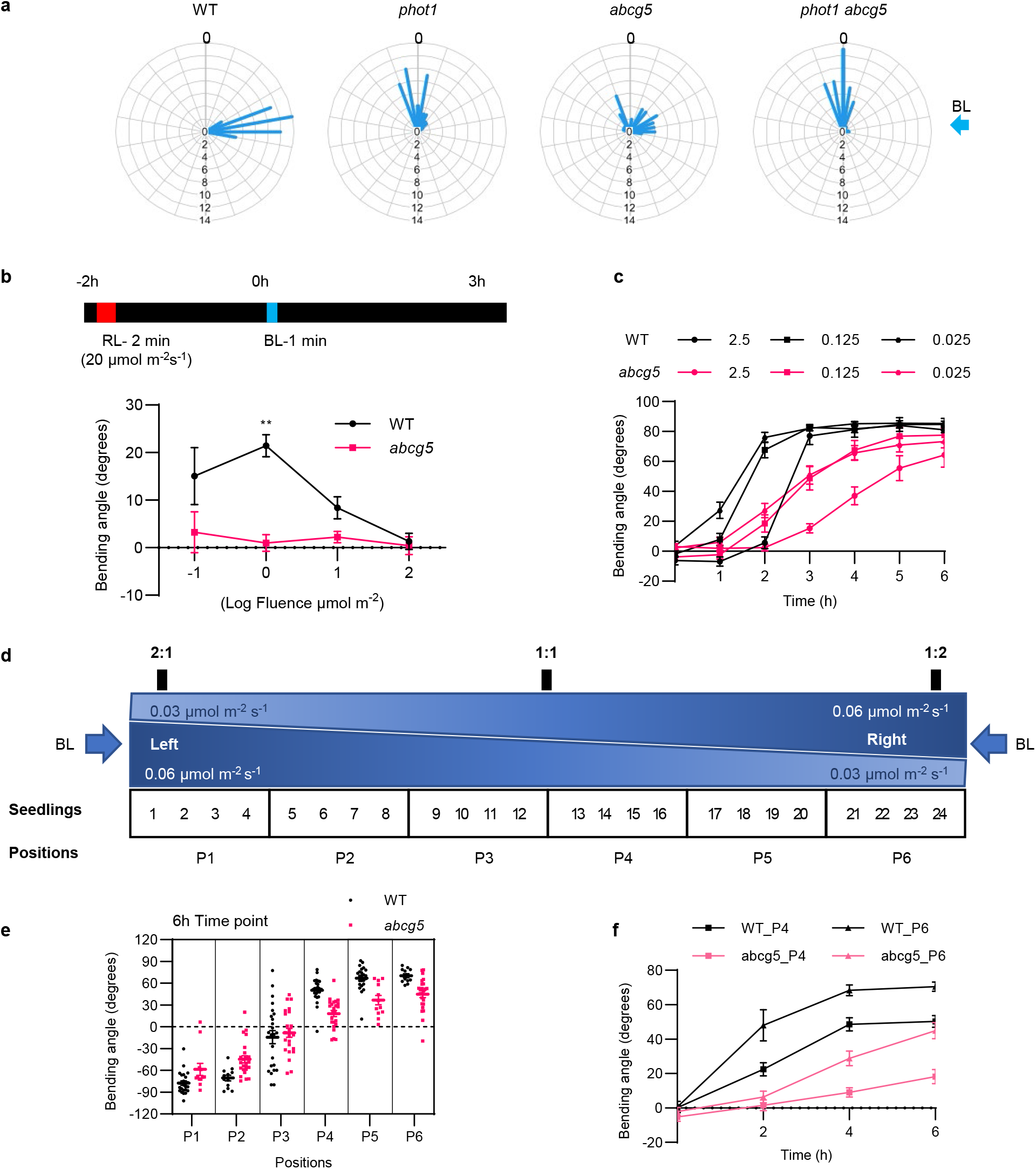
Random bending of *abcg5* is phot1-dependent. (a) Radar plots showing the results phototropic assay. Three-day-old etiolated seedlings grown on vertical plates were treated with unilateral light (0.025 μmol m^-2^ s^-1^) for 24 h and measured final angle relative to vertical. The bar length in the radar represents the relative frequencies of hypocotyl angles at 10 ° intervals; n=43-46. (b) Pulse-induced first positive response. Top: Experimental scheme. Three-day-old etiolated seedlings grown on vertical PCR tubes stimulated with red light (20 μmol m^-2^ s^-1^) for 2 min. Two hours later they were stimulated with a 1-min BL of various fluences. Bottom: Data are mean ± SEM of the final growth angle relative to vertical, pulled from independent experiments, each with n=11-12. **P<0.01 in ANOVA followed by Šídák’s multiple comparisons test. (c) Time-course analysis of a phototropic assay. Three-day-old etiolated seedlings grown on al PCR tubes illuminated from one side with continuous fluence rates of BL (0.025, 0.125, and 2.5 μmol m^-2^ or six hours and measured the final growth angle relative to vertical (mean ± SEM;n=24). (d) Experimental me for bidirectional illumination of seedlings. Three-day-old etiolated seedlings grown on 24 PCR tubes at positions P1-P6 were exposed to 0.06 μmol m^-2^ s^-1^ blue light from both sides creating a shallow light gradient at position P3 and P4 and a steep gradient at position P1, P2, P5, P6. (e) Bending angles of WT and *abcg5* at different positions (P1-P6). The positive angle represents bending toward the right side, while the negative s show bending toward the left side. (f) Kinetics of bending of seedlings at position P6 and P4. Data are ± SEM; pulled from three independent experiments, each with n=11-16.

**Extended data Figure 4.**
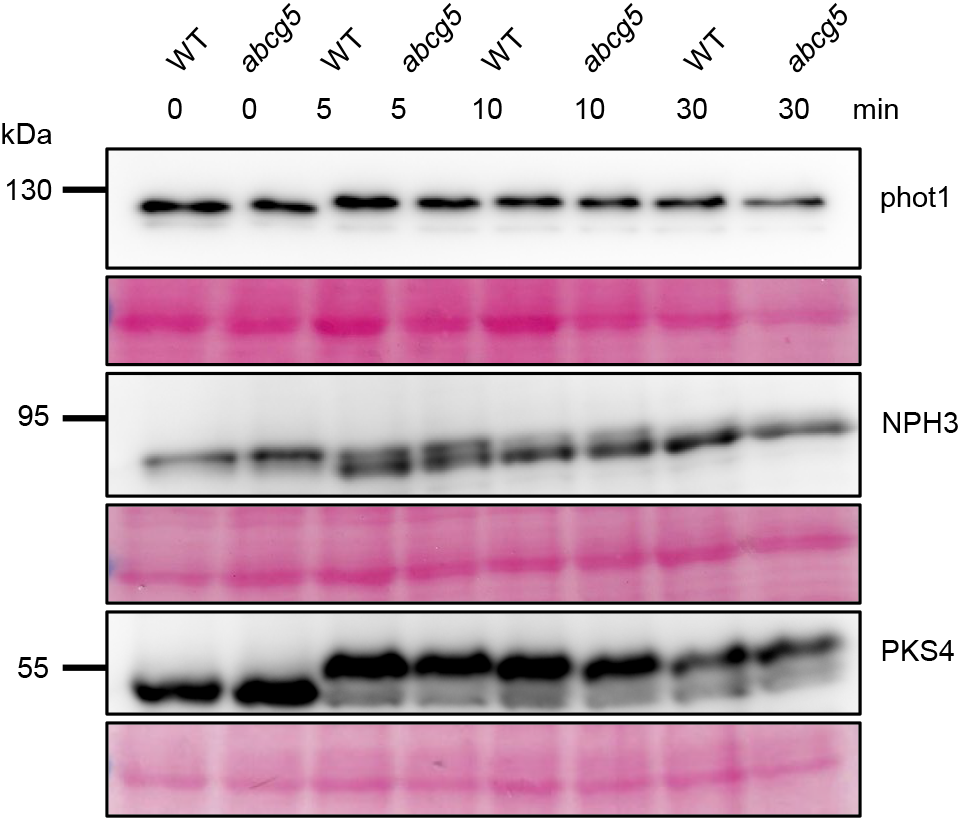
*abcg5* showed normal phot1 signaling. The phosphorylation status of phot1, NPH3, PKS4 in 3-day-old etiolated seedlings of WT and *abcg5* after illuminating with blue light (2.5 μmol m^-2^ s^-1^). proteins were extracted at the indicated times, separated by SDS–PAGE, transferred onto nitrocellulose membrane, and probed with antibodies against phot1, NPH3, and PKS4. Ponceau red staining is shown for each membrane as loading control.

**Extended data Figure 5.**
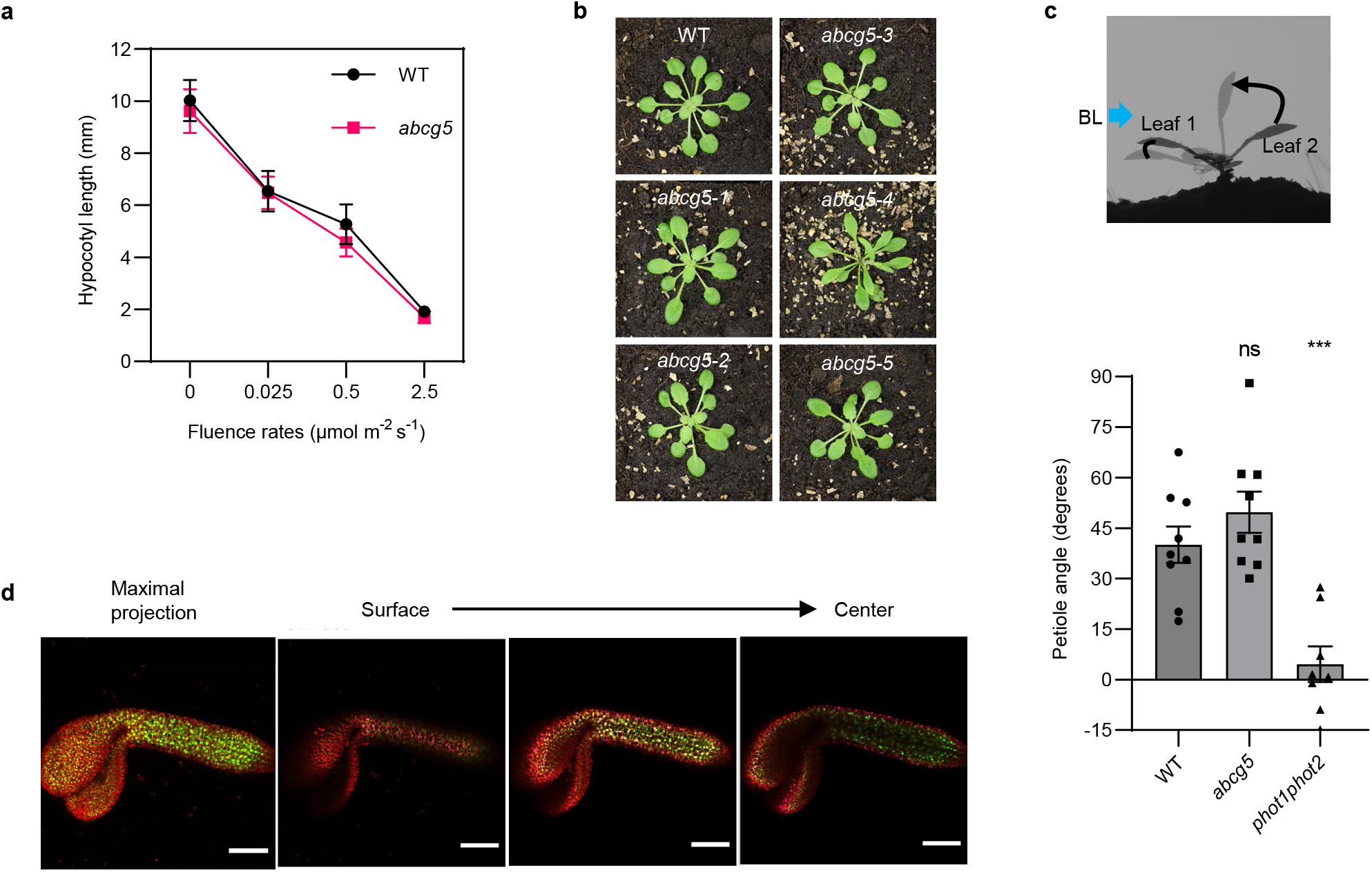
ABCG5 function is dispensable for de-etiolation and leaf phototropism. (a) Hypocotyl lengths of 3-day-old seedlings grown under continuous BL with fluence rates of 0.025 to 2.5 μmol m^-2^ (mean ± SEM; n=53-61 seedlings). (b) Pictures of 4-week-old WT and *abcg5* plants grown in long days (LD, ours light, 8 hours dark). (c) Phototropic response of leaf 1 and 2 of two-week-old plants grown in LD irradiated with unilateral BL (1 μmol m^-2^ s^-1^) for 24 h. The leaf angles were measured relative to the horizontal. In response to blue light, the angle of leaf 1 decreases while the angle of leaf 2 increases. The change in angle for and 2 was plotted on the bar graph. The data represented is a mean ± SEM, pulled from two independent experiments and a total n=8-9. Statistics by ANOVA followed by Dunnett’s multiple comparisons test are shown; * * * P<0.001 and NS, no significance. (d) NLS-3X Venus under *ABCG5* promoter. Maximal projection image obtained by using Z-stacks from surface to center of the embryo. The red color shows the chlorophyll autofluorescence while green color denotes Venus signal.

**Extended data Figure 6.**
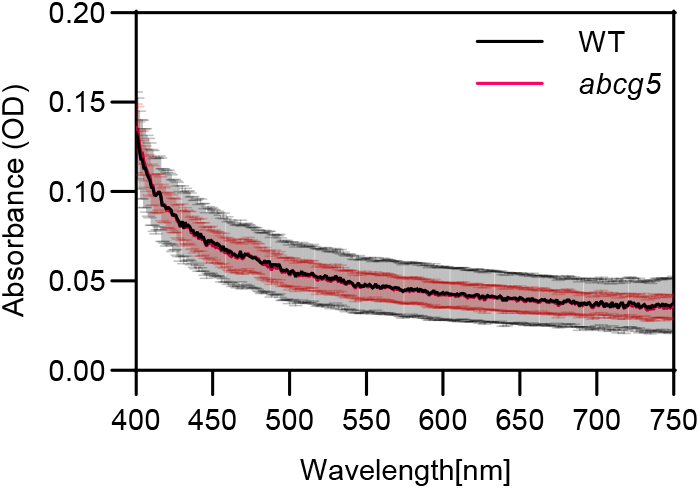
No difference between WT and *abcg5* light absorption properties. Absorption spectra of crude extracts from ∼100 dark-grown seedlings of WT and *abcg5*. For crude extracts, samples were crushed in an Eppendorf tube using pestles and centrifuged at 13000 rpm for 5 min. The soluble crude on of 30 μl was collected in a fresh tube, and 20 μl of sterile water was added to make the final 50 μl volume. The absorption scan for the visible spectrum (400-750 nm) of crude extracts were measured using a n Safire 2 plate reader. The data represented is mean ± SEM;n=5

**Extended data Figure 7.**
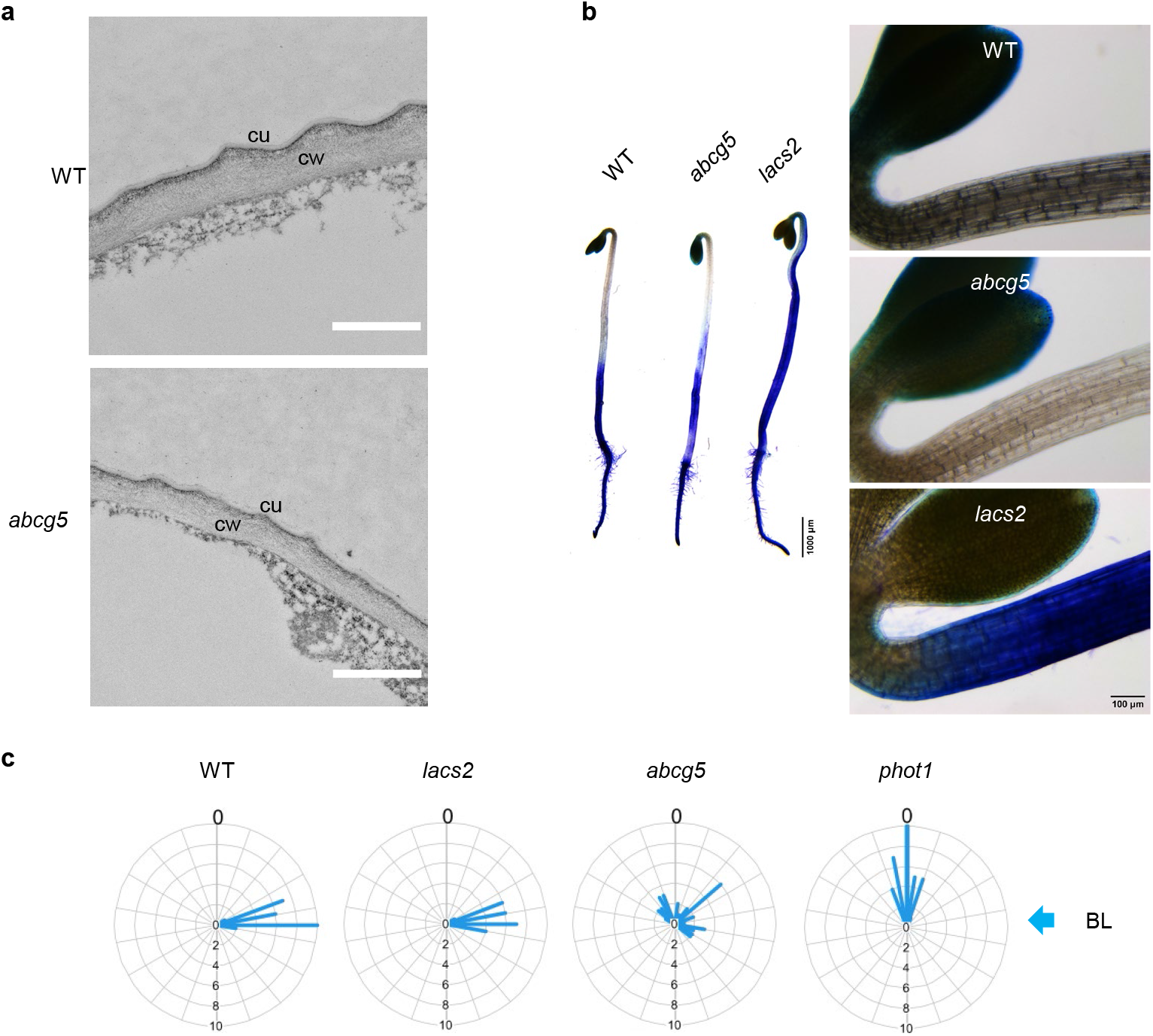
Cuticular defects are not responsible for phototropic defects. (a) Transmission on microscopy (TEM) images of transverse cuts of a 3-day-old etiolated seedlings hypocotyl. cu, cuticle; ell wall. Scale bar 1 μm. (b) Representative pictures of toluidine blue (TB) staining assay of 3-day-old ted seedlings of WT, *abcg5*, and *lacs2*. Plant materials were incubated for 10 min at room temperature in a 0.05 % TB in 0.4% Tween solution, followed by a gentle wash with water to remove excess TB (c) Radar plots showing hypocotyl bending in a phototropic assay. Three-day-old etiolated seedlings grown on vertical plates treated with unilateral blue light (0.025 μmol m^-2^ s^-1^) for 24 h and measured final growth angle relative to al. The bar length represents the relative frequencies of hypocotyl angles in a 10 ° interval. n=25-37.

**Extended data Figure 8.**
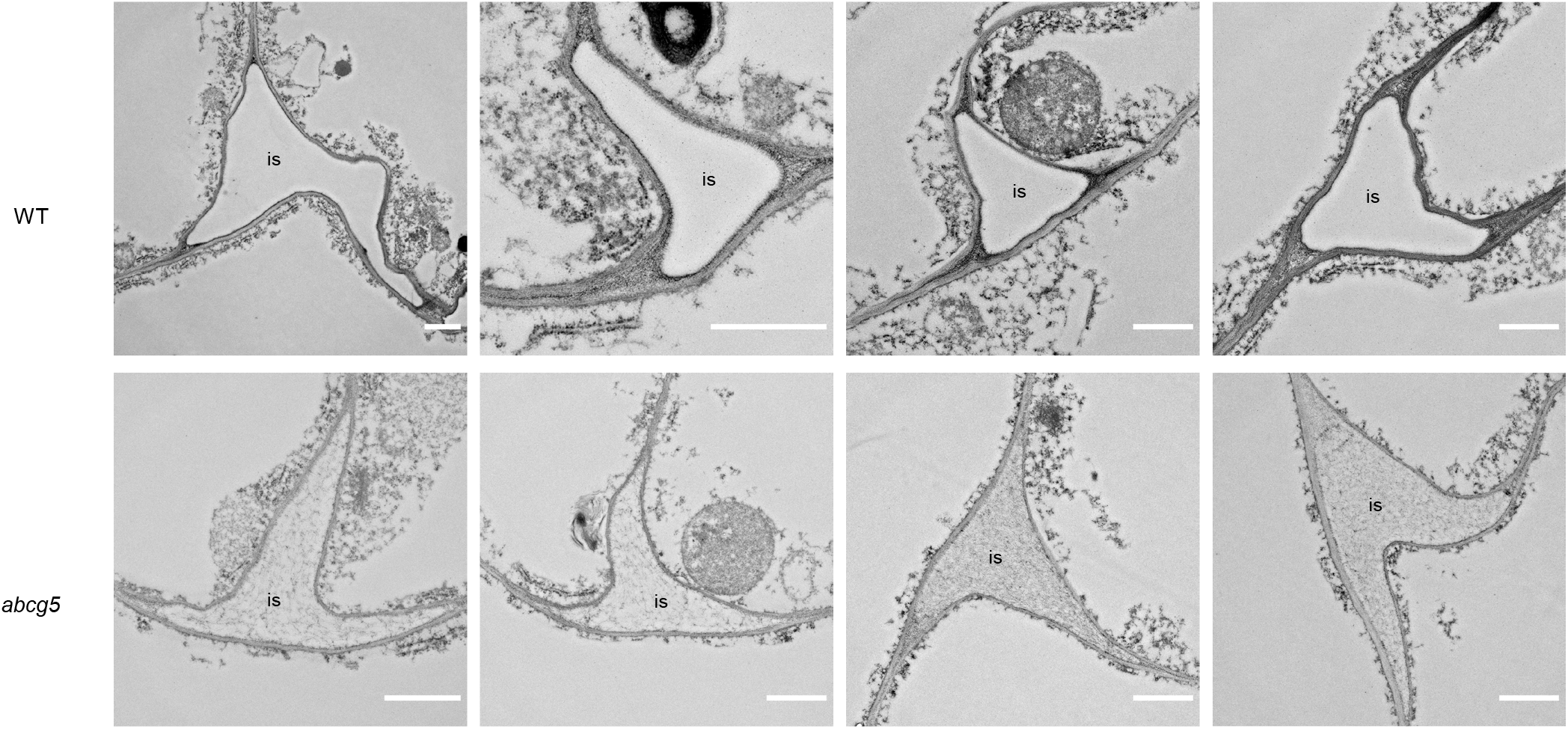
Differences between WT and *abcg5* intercellular spaces shown by TEM. Transmission electron microscopy (TEM) images of transverse cuts of a 3-day-old etiolated seedlings’ hypocotyls. is, intercellular space. Scale bar 1μm.

**Extended data Figure 9.**
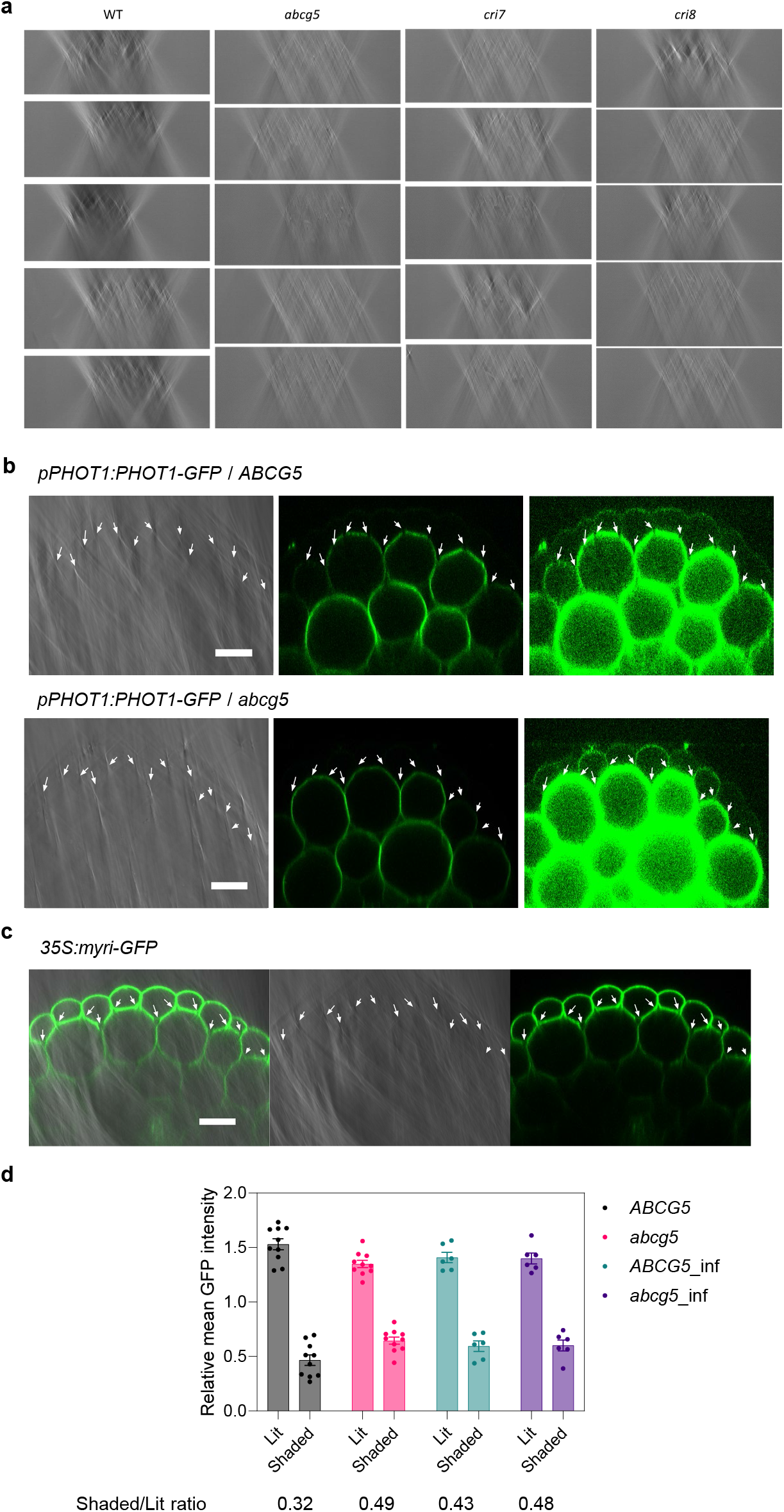
Air bubbles create light signals within the tissue. (a) xz orthogonal view of a bright z-stack of etiolated hypocotyls. Five independent plants are shown for each genotype. (b) xz orthogonal of bright field and fluorescence confocal images of etiolated hypocotyls expressing *pPHOT1:PHOT1-GFP* ABCG5 wild tipe or mutant (*abcg5*) backgrounds. Left: bright field, Middle: fluorescence, Right: fluorescence with enhanced brightness. Arrows show the intercellular spaces between epidermal and cortex Scale bar 20 μm. (c) xz orthogonal view of bright field and fluorescence confocal images of etiolated hypocotyls of plants expressing *35S:myri-GFP*. Left: merge, Middle: bright field, Right: fluorescence. Arrows the intercellular spaces between epidermal and cortex cells. Scale bar 20 μm. (d) Quantification of fluorescence confocal images of plants expressing *pPHOT1:PHOT1-GFP* in an ABCG5 wild type (*ABCG5*) or nt (*abcg5*) backgrounds, live or vacuum infiltrated with 4% PFA (*ABCG5*_inf, abcg5_INF). xz orthogonal of xz scans were quantified. For each image, the mean GFP fluorescence of the whole transverse cut, as well as of the half closer to the objective (lit) and farther from the objective (shaded) were quantified. Plotted s are mean GFP fluorescence of each side relative to the mean GFP fluorescence of the whole image. The shaded/lit ratio is calculated from these relative values.

## Notes

### Competing Interest Statement

The authors have declared no competing interest.

