## Supplementary material for "Air channels create a directional light signal to regulate hypocotyl phototropism": Materials – Methods

### Supplementary Methods

#### Plant materials

All Arabidopsis lines used in this study are derived from the Columbia (*Col*) accession. The T-DNA insertion lines *abcg5-1* (GABI_726G11), *abcg5-2* (GABI_367D06)[1], *abcg5-3* (SALK_074250), and *abcg5-4* (SAIL_700_E01) for ABCG5 were obtained from NASC stock center. Most of the physiological assays were performed using this *abcg5-5* (*abcg5*) allele (see below). The hyperhydric *cristal* (*cri7* and *cri8*) mutants were previously described[2]. Although it is known that the *cri7* and *cri8* phenotypes are due to monogenic recessive mutations, the underlying genes are not characterized. By sanger sequencing, we confirmed that *cri7* and *cri8* have a wild-type ABCG5 allele. The previously reported mutants *phot1-5*, *phot1-5 phot2-1*[3]; *pin3-3 pin4-101 pin7-101*[4] and *lacs2-3*[5] were used. The double mutant *phot1-5 abcg5-5* was generated by crossing. We used the pPHOT1: PHOT1-GFP in *phot1-5 phot2-1* background [6] line to obtain the pPHOT1: PHOT1-GFP in *phot1-5 phot2-1 abcg5-5* background by genetic crossing.

#### Identification of causal mutation

The point mutant *abcg5-5* (G538R) was isolated from an EMS mutagenized genetic screen to identify mutants defective in long-term phototropic response. In brief, we screened 12-18 M2 seedlings from every 2250 M2 families. The identified mutant lines were repeatedly backcrossed to WT plants, resulting in a segregating population pooling into two groups- mutant and WT pool. The gene mutation responsible for the underlying phenotype was identified by whole genome sequencing (WGS) comparison between these two pools. Briefly, paired WGS-reads were trimmed to remove adapters (trimmomatic v0.36; ILLUMINACLIP:config/TruSeq3-PE.fa:2:30:10 MINLEN:50) and aligned against the TAIR10 reference genome (bwa mem v0.7.15, samtools v1.4). After de-duplication (picard -tools 2.9.0; SortSam, MarkDuplicates, MergeSamFiles, AddOrReplaceReadGroups: VALIDATION_STRINGENCY=LENIENT; ) and interval-realignment ( GATK v3.4-46; RealignerTargetCreator + IndelRealigner), SNVs were called for both genotypes (GATK v3.4-46; UnifiedGenotyper ; -glm BOTH). The VCFs with the variants were compared in a custom script, to identify novel, homozygous variants in the final line. Therefore, we filtered variants based on quality (30), minimum variant allele frequency (VAF) of 50% , coverage of 50x and a maximum 20% VAF for the reference sample. The potential effect of the variant on coding genes was analyzed using snpEff (snpEff v3.6c; athalianaTair10), which led to the identification of the mutated ABCG5 gene. The identified point mutation was GGT-CGT.

#### Materials generated in this study

For cloning purposes, we used pFR100/101 vectors derived by replacing the original seed coat selection marker [7] with a FAST-red (pOLE1: OLE1-TagRFP) seed coat selection marker[8]. We amplified the *ABCG5* gene (1947 bp) from genomic DNA using AG429/AG430 primers and GFP-S65T sequence from plasmid pCF203 using AG427/AG428 primers. These two fragments were cloned into pFR101 to get p35S: GFP-ABCG5 (pGMN01) construct using an infusion cloning kit. We amplified the ABCG5 promoter (2229 bp) and ABCG5+3' UTR (2188 bp) from genomic DNA using primer pairs AG406/AG431 and AG429/AG407, respectively. The GFP-S65T sequence was amplified from plasmid pCF203 using AG432/AG428 primers. All three amplified fragments were cloned into pFR100 to get pABCG5: GFP-ABCG5 (pGMN02) construct. To generate the pABCG5 reporter construct (pGMN09), we digested pGMN02 plasmid with HindIII to release the GFP-ABCG5-3' UTR sequence and replaced it with NLS-3X mVenus sequence amplified from pSF376 using GMN101/102 primers. All constructs were transformed into pSoup containing Agrobacterium strain GV3101. The constructs pGMN01/02 were transformed into the *abcg5-5* background and pGMN09 in the WT background using the floral dip method described previously [9]. Single insertion homozygous T3 lines were selected based on the seed coat selection marker.

**Plant growth conditions**

For physiological experiments, seeds were surface-sterilized using 70% ethanol and 0.05% Triton-X for 5 min and a second wash with 100% ethanol for 5 min. Seeds were sown on square Petri dishes containing ½ MS (0.8% Agar) or sterile wet filter papers. Seeds were stratified at 4 °C in the dark for 3-4 days. We induced the germination by 4-6 hours of white light (80 μmol $m^{-2}$ $s^{-1}$) treatment and kept the plate back in the dark for 3-days at 19 °C or at different blue light intensities (0.025 to 2.5 μmol $m^{-2}$ $s^{-1}$) for analyzing de-etiolation of seedlings. For PCR tube assay, stratified seeds after induction of germination were placed on the 1/2MS (0.8% Agar) in PCR tubes (modified from [10]). PCR tubes were placed in the box with wet filter papers inside to avoid the desiccation of the media. These boxes were covered with aluminum foil and kept in the dark for 3-days at 19 °C. Stratified seeds from filter paper were placed on soil pots and grown at 22 °C on a long day (16h Light: 8h Dark).

**Physiological experiments**

For phototropic stimulation, the 3-day-old etiolated seedlings grown on vertical plates were treated with unilateral blue light (BL) (0.025 μmol $m^{-2}$ $s^{-1}$) for 24 hours, while for gravi-stimulation, plates were rotated by 90 ° and pictures were captured using infra-red CCD camera system at different time points during 24h. The angles formed by the hypocotyls relative to vertical (after phototropic stimulation) or horizontal (after gravi stimulation) were measured using ImageJ. Seedlings were classified into 10 ° angle intervals (-10 ° – 0 °, 0 ° – +10 °, +10 ° – +20 °…… -20 ° – -10 °) and the radar plots were drawn using RStudio.

For pulse-induced first positive phototropic response, 3-day-old etiolated seedlings growing on individual PCR tubes were pretreated with an overhead red light (RL) at 20 μmol $m^{-2}$ $s^{-1}$ for 2 min. The seedlings were arranged on custom-made PCR tube stands (24 seedlings per row) so that the directional BL would be perpendicular to the plane of the hook (to avoid differences in the response due to irradiating the hook or cotyledons). Two hours after the RL pulse, a BL pulse of various intensities (1.7, 0.17, 0.017, and 0.0017 μmol $m^{-2}$ $s^{-1}$) was applied for 1 min. The pictures were captured before and 3 hours after the BL pulse using an infra-red CCD camera system.

For continuous BL light-induced phototropic response, the seedlings arranged on the PCR tubes stand were illuminated from one side with continuous fluences of BL (0.025, 0.125, and 2.5 μmol $m^{-2}$ $s^{-1}$). For the shallow light gradient experiment, etiolated seedlings arranged on PCR tube stands at positions P1-P6 were exposed to 0.06 μmol $m^{-2}$ $s^{-1}$ blue light from both sides creating a shallow light gradient at positions P3 and P4 and a steeper gradient at position P1, P2, P5, P6. The pictures were captured every 10 min using a photo camera equipped with an automatic trigger.

**Hypocotyl transparency analysis**

The images of the upper hypocotyl region of 3-day-old etiolated seedlings were captured using a Leica M205 FCA stereomicroscope. ImageJ was used to select the circular regions of interest (ROI). The percent transmitted light was calculated as a ratio of the gray values from a ROI on the hypocotyl divided by the mean gray values from a ROI on an empty region. Per genotype, 8-9 seedlings were analyzed and plotted as a bar graph using GraphPad.

#### Leaf angle measurement

We used two-week-old soil-grown seedlings with fully developed leaves 1 and 2 (~6-8mm length) to analyze the green tissue's phototropic response. At ZT0, the seedlings were illuminated with directional BL (1 µmol $m^{-2}$ $s^{-1}$) parallel to the plane of leaves 1 and 2. The pictures were captured using an infra-red CCD camera system before BL illumination and after 24 hours of BL exposure. We measured the leaf angles relative to the horizontal. In response to blue light, the angle of leaf 1 reduces while leaf 2 angle increases. Total leaf angle change was calculated by the addition of angle change for leaf 1 and leaf 2 and plotted as a bar graph using GraphPad.

#### Toluidine blue staining assay

We assessed the cuticular integrity using a toluidine blue (TB) staining assay as previously described [11]. Three-day-old etiolated seedlings were incubated for 10 min at room temperature in a 0.05% TB in 0.4% Tween solution, followed by a gentle wash with water to remove excess TB. The pictures were captured using a Leica M205 FCA stereomicroscope.

#### Western blot

Seedlings were grown on horizontally orientated plates for 3 days in darkness at 22 °C and illuminated with BL (2.5 µmol $m^{-2}$ $s^{-1}$) for 0 – 30 min. Total proteins were extracted by grinding ~50 seedlings in 50 µl 2X Laemmli buffer with 10% β-mercaptoethanol. Samples were heated for 5 minutes at 95 °C and centrifuged for 5 minutes; 10 µl per sample were loaded on 8% SDS/PAGE gels. Electrophoresis was performed at 100V for 10 min and 200V for 40 min. The transfer was performed using the TransBlot Turbo system at 25V for 7 min (BioRad). Detection of endogenous phot1, NPH3, and PKS4 was done using anti-phot1 (1:10,000) [12], anti-NPH3 (1:3,000) [13], and anti-PKS4 (1:300) [14] in PBS 5% milk 0.1% tween (PBSTM). Membranes were blocked for 1 h at room temperature with PBSTM and incubated overnight with the corresponding primary antibody at 4 °C. Chemiluminescence signals were generated using Immobilon Western HRP Substrate (Millipore). Signals were detected with a Fujifilm ImageQuant LAS 4000 mini-CCD camera system.

#### Analysis of absorption spectra of a soluble fraction from crude extracts

Seedlings were grown on horizontally orientated plates for 3 days in darkness at 22 °C, and ~100 seedlings were crushed in an Eppendorf tube using pestles and centrifuged at 13000 rpm for 5 min. The soluble crude fraction of 30 µl was collected in a fresh tube, and 20 µl of sterile water was added to make the final 50 µl volume. The sterile water was used as a blank. We took a total of 50 µl of the sample into the transparent flat bottom 96 well plates for absorption analysis. The absorption scan for the soluble crude extracts' visible spectrum (400-750 nm) was measured using the Tecan Safire 2 plate reader. Five biological replicates were used, and the experiment was repeated two times.

#### Floating assays

For the floating assay, we either used intact seedlings or dissected roots, hypocotyls and cotyledons from the etiolated seedlings grown on horizontally orientated plates for 3 days in darkness. We quantified the difference in buoyancy by placing ~10 samples in 25 ml beakers filled with sterile water and incubating for 10 mins at room temperature. The number of floating samples was counted and expressed as a percent floating sample. Four biological replicates were used, and the experiment was repeated two times.

#### Cryo-scanning electron microscopy (Cryo-SEM)

For cryo-scanning electron microscopy, we used a Quorum system PP3000T (Quorum Technologies) attached to a Quanta 250 FEI scanning electron microscope (FEI Company). Three-day-old etiolated seedlings were mounted on aluminum stubs using a mixture of Tissue-Tek and colloidal graphite (50:50), frozen in nitrogen slush at -225 °C for two times and then transferred to the preparation chamber of the Quorum system. We set the Quorum chamber temperature at -175°C, and to avoid contamination; vacuum pressure was set at 6e-6 bar. Seedlings in the Quorum-chamber were fractured at the upper hypocotyl region and then sputter-coated with platinum at 10 mA for 60 s. After transfer on the cryo-stage at -140 °C in the scanning electron microscope, imaging was performed, on the transverse hypocotyl cuts, at 7 keV using backscattered detector. For each genotype, minimum 5 hypocotyls were scanned. The images were analyzed using ImageJ to detect air spaces at the intercellular spaces.

#### 3D Non-Destructive X-Ray Analysis

All samples were measured with an X-ray micro-CT scanner (Ultratom, RX-Solutions). The scanning protocol and parameters were kept identical for all seedling samples. A Hamamatsu microfocus X-ray sourcce (160 kV) was used, in transmission mode, with a 0.1 mm thick aluminium filter, a LaB6 cathode and Tungsten target. The acquisition was performed with a current of 80 mA and a voltage of 40 kV. During volume data acquisition, the sample rotated by 360 degrees and 800 projections were taken by step of 0.45 degree to ensure a very precise volume reconstruction. The X-ray beam attenuation was registered by a Varex Paxscan 2520 plane detector of 1920 x 1536 pixels, with an exposure time of 0.33 s. The projections were then processed (X-act RX-Solutions, Filtered Backprojection) to reconstruct a corrected volume composed of around 1500 slices in 16 bit Tiff format. Voxels dimensions were typically of 1 x 1 x 1 microns. Each scan lasted 18 minutes. To keep the sample still and avoid dehydration during image acquisition, hypocotyls were covered with a layer of all-purpose transparent glue (UHU) and glued to a P10 plastic tip, which was later positioned on the sample holder.

### Transmission Electron Microscopy

Arabidopsis hypocotyls were fixed in 2.5% glutaraldehyde solution (EMS, Hatfield, PA) in phosphate buffer (PB 0.1 M [pH 7.4]) for 1 hour at room temperature and subsequently fixed in a fresh mixture of osmium tetroxide 1% (EMS) with 1.5% potassium ferrocyanide (Sigma, St. Louis, MO) in PB buffer for 1 hour at room temperature. The samples were then washed twice in distilled water and dehydrated in ethanol solution (Sigma, St Louis, MO, US) in a concentration gradient (30% for 40 minutes; 50% for 40 minutes; 70% for 40 min and 100% for 1 hour 3 times. This was followed by infiltration in LR White resin (EMS, Hatfield, PA, US) in a concentration gradient (33% LR White 33% in ethanol for 6 hours; 66% LR White in ethanol for 6 hours; 100% LR White for 12 hours two times) and finally polymerized for 48 hours at 60°C in an oven in atmospheric nitrogen. Ultrathin sections (50 nm) were cut transversely to the hypocotyl, using a Leica Ultracut UC7 (Leica Mikrosysteme GmbH, Vienna, Austria), picked up on a copper slot grid 2x1mm (EMS, Hatfield, PA, US) and coated with a polystyrene film (Sigma, St Louis, MO, US). Micrographs and panoramic images were taken with a transmission electron microscope FEI CM100 (FEI, Eindhoven, The Netherlands) at an acceleration voltage of 80kV with a TVIPS TemCamF416 digital camera (TVIPS GmbH, Gauting, Germany) using the software EM-MENU 4.0 (TVIPS GmbH, Gauting, Germany).

### Confocal microscopy

To image embryos expressing pABCG5:NLS-3xVenus an LSM710 inverted confocal microscope equipped with an EC Plan-Neofluar 20x/0.50 M27 objective (Zeiss) was used. Samples were excited with an 514nm laser and detection was performed at 519-557nm for Venus (green), 593-616nm for TagRFP (magenta) and 632-687nm for Chlorophyll (red). A z-stack was performed and representative images of the upper, middle and center of the embryo are shown.

For images in Fig. 3b and Extended data Fig. 9 an LSM880 inverted confocal microscope equipped with a Plan-Apochromat 20x/0.8 M27 objective was used. For the close-up looks in Figure 3c and

Extended data Fig. 9b, c, a Plan-Apochromat 63x/1.4 Oil DIC M27 objective was used. Samples were excited with a 488nm laser and detection was performed at 496-531nm for GFP and with the T-PMT for bright field. In all cases a Z-stack was performed and the XZ orthogonal view is shown.

For the quantifications in Fig. 3d and Extended data Fig. 9d a DMi 8 Stellaris inverted microscope equipped with a HC PL APO CS2 20x/0.75 DRY objective (Leica) was used. Samples were excited with a 488nm laser and detection was performed at 505-530nm with a HyD S detector in photon counting mode. XZ scans were performed. ImageJ was used for quantification.

**Optical properties analysis using the integrating sphere**

For optical property analyses, an integrating sphere (RT-060-SF, Labsphere) was used to measure transmittance and/or reflectance. A solar simulator Mercury-Xenon lamp (Thermo Oriel, model 66902) at 300W was used as a light source to illuminate the samples. A spectrometer (Oriel, model 77400, MultiSpec 125TM, type 1/8m) was used for measurements.
3-day-old etiolated seedlings were grown on horizontally orientated plates. Cotyledons were discarded and only hypocotyls and roots were used for the analysis. We used vacuum treatment to infiltrate the hypocotyl samples with 0.025% Silwet L-77 solution. The hypocotyl samples with and without infiltration were mounted in a chamber prepared by sticking two SecureSeal$^{TM}$ imaging spacers (20 mm DIA x 0.12 mm Depth) on a microscope glass slide and filled with a 0.025% Silwet L-77 solution in water. The samples were covered using a glass coverslip. The mock samples were prepared using Silwet L-77 solution without any hypocotyls.
To measure transmittance, samples were placed at opening 1 while covering the openings 2 and 3 using white board (Supplementary Fig. 1). For diffused transmittance, opening 2 was opened allowing unscattered light to escape the sphere. For reflectance measurements samples were placed at the opening 2 and leaving opening 1 open and opening 3 closed. For diffused reflectance, the sample was placed at opening 2 and opening 3 was open allowing unscattered reflected light escaping the integrating sphere (Supplementary Fig. 1).
5-8 independent biological samples were measured. For each sample, the average of three or two technical replicates was measured for transmittance and reflectance, respectively. The same sample was used to measure total and diffused transmittance or total and diffused reflectance. Two experiments for transmittance and two experiments for reflectance were done.

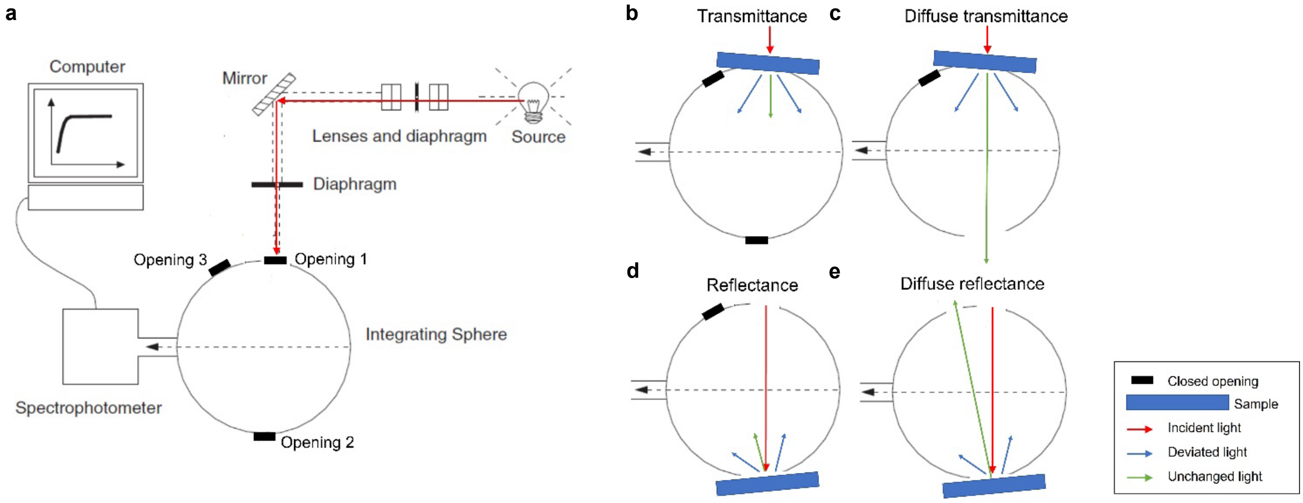


**Supplementary Figure 1. Measurements with the integrating sphere.** (a) Diagram of the integrating sphere and its components. (b-e) – Setup to measure transmittance (b), diffuse transmittance (c), reflectance (d) and diffuse reflectance (e).

**Supplementary Table 1: Genotyping conditions to select mutations in crosses**

| No. | Allele | PCR conditions | Digestion enzyme | Result |
|---|---|---|---|---|
| 1 | *abcg5-1* | AG415+AG414, 55°C<br>AG415+AG156, 55°C | | WT: 1021 bp<br>Mutant: ~700 bp |
| 2 | *abcg5-2* | AG413+AG412, 55°C<br>AG413+AG156, 55°C | | WT: 1193 bp<br>Mutant: ~700 bp |
| 3 | *abcg5-3* | AG408+AG409, 55°C<br>AG409+GMN063, 55°C | | WT: 1192 bp<br>Mutant: ~700 bp |
| 4 | *abcg5-4* | AG410+AG411, 55°C<br>AG411+GMN064, 55°C | | WT: 957 bp<br>Mutant: ~600 bp |
| 5 | *abcg5-5* | AG392+AG393, 55°C | AgeI | WT: 186+ 24 bp<br>Mutant: 210 bp |
| 6 | *phot1-5* | CF342 + CF343 + CF344 + CF345, 55°C | | WT:250 + 450 bp<br>Mutant:250 bp |
| 7 | *phot2-1* | CF346 + CF347, 55°C | Mbo1 | WT:398 +230+120 bp<br>Mutant: 530+230 bp |

**Supplementary Table 2: Primers used in this study**

| No. | Primer Name | Sequence (5'-3') | Purpose |
|---|---|---|---|
| 1 | AG156 | CCCATTTGGACGTGAATGTAGACAC | Genotyping |
| 2 | AG392 | TCATAGTTGGAAACTCAGTGATTACC | Genotyping |
| 3 | AG393 | CAGTCACCAAACATTTTCCGAATCC | Genotyping |
| 4 | AG406 | AATCCAGTGGGTACCCGGGGATCCTTTTTCCTAAGAAAAAAACTTGAAAT | Cloning |
| 5 | AG407 | TTGCATGCCTGCAGGTCGACTCTAGATGTAAGTAACTTTGCTGGAATTG | Cloning |
| 6 | AG408 | TTGGGTCAAAGATTGCTTTTGTAG | Genotyping |
| 7 | AG409 | GTTACCGGTTTCATCCTCTAGCTTC | Genotyping |
| 8 | AG410 | GCACTGAACAGAAGGGTTTCCTCC | Genotyping |
| 9 | AG411 | CATGTAAATCGATTTGCCGAAAAG | Genotyping |
| 10 | AG412 | GCAAAGAGCATGATTGAGGAGATG | Genotyping |
| 11 | AG413 | TCTCAACCCCAGTGTACTGGCTGG | Genotyping |
| 12 | AG414 | GTGAATTTGCCACTTTTGCTCTCC | Genotyping |
| 13 | AG415 | CATCAATGGAGAAGCAAGGATGTG | Genotyping |
| 14 | AG427 | GGACACGCTGACAAGCTGACTCTAGAATGAGTAAAGGAGAAGAACTTTTC | Cloning |
| 15 | AG428 | ACATCCTTGCTTCTCTTTGTATAGTTCATCCATGCC | Cloning |
| 16 | AG429 | TGAACTATACAAAGAGAAGCAAGGATGTGAGATC | Cloning |
| 17 | AG431 | TTCTCCTTTACTCATTGATGAAGAAGCTTTGTTCTC | Cloning |
| 18 | AG432 | AGCTTCTTCATCAATGAGTAAAGGAGAAGAACTT | Cloning |
| 19 | GMN063 | ATTTTGCCGATTTCGGAAC | Genotyping |
| 20 | GMN064 | TAGCATCTGAATTTCATAACCAATCTCGATACAC | Genotyping |
| 21 | GMN101 | AGAGAGAACAAAGCTTCTTCATCAGAATTCATGCCAAAGAAGAAGAG | Cloning |
| 22 | GMN102 | AGCTCAAGCTAAGCTAGGTCACTGGATTTTGGTTTTAGGAATTAGAAA | Cloning |
| 23 | CF342 | TGTTGGCATCAGGAAGTT | Genotyping |
| 24 | CF343 | TGTGGCAGGAAAGAAGTT | Genotyping |
| 25 | CF344 | TGCCTGCAAACCAATAAC | Genotyping |
| 26 | CF345 | CCGGAGCAGGACATACG | Genotyping |
| 27 | CF346 | CTGCCTCACAATAAGGAGAG | Genotyping |
| 28 | CF347 | GAACCTTGCAGAGTCTTCTG | Genotyping |
